# An unusually long 5’UTR of ATP dependent cold shock DEAD-box RNA helicase gene *csdA* negatively regulates its own expression in *Escherichia coli*

**DOI:** 10.1101/2021.03.08.434349

**Authors:** Soma Jana, Partha P. Datta

## Abstract

<u>C</u>old-<u>s</u>hock <u>D</u>EAD-box protein <u>A</u> (CsdA) is an ATP dependant cold shock DEAD-box RNA helicase. It is a major cold shock protein needed for the cold adaptation in *Escherichia coli*. Although the CsdA has been studied at the protein level, further studies are necessary to understand its mechanisms of gene regulations. In this regard, we have constructed a promoter less vector with the ORF of a GFP reporter and found that the promoter of the *csdA* gene resides far upstream (more than 800 bases) of its coding region. Furthermore, our *in vivo* deletion experiment has confirmed the existence of this extraordinarily long 5’UTR. Our results show that it represses its own expression. In addition, the short peptide encoding (26 aa) *yrbN* gene resides within this 5’UTR as an operon with 8 overlapping nucleotides with the *csdA* coding region. Besides, we observed that the *csdA* gene expression may also occur along with immediate upstream (180 nucleotides) *nlpI* gene both at 37°C and 15°C and from the *pnp* gene (1173 nucleotides upstream) only during cold. In conclusion, *csdA* gene has operon feature like prokaryotes, in contrast, it also contains an extraordinarily long 5’UTR, found in eukaryotes.

## Introduction

Bacteria produce cold shock proteins, when the culture is shifted to low temperature. Cold shock response has been studied in many bacteria like, mesophilic bacteria; *Enterococcus faecalis* (Panoff *et al*., 1997), *Bacillus subtilis* (Lottering and Streips, 1995; Graumann *et al*., 1996), and *Lactococcus lactis* (Panoff *et al*., 1994), psychrotrophic bacteria; *Arthrobacter globiformis* (Berger *et al*., 1996), *Pseudomonas fragi* (Hebraud *et al*., 1994; Michel *et al*., 1996; 1997), *Bacillus psychrophilus* (Whyte and Inniss, 1992), *Pseudomonas putida* (Gumley and Inniss, 1996), *Vibrio vulnificus* (McGovern and Oliver, 1995), *Rhizobium* (Cloutier *et al*., 1992), and *Listeria monocytogenes* (Phan-Thanh and Gormon, 1995; Bayles *et al*., 1996), and psychrophilic bacterium; *Aquaspirillum arcticum* (Roberts and Inniss, 1992). During this cold shock condition, the secondary structure of the nucleic acids stabilizes that causes reduction of the rate of DNA replication, transcription, and translation processes (Phadtare, 2004). Cold shock proteins play important roles at this condition to resume the cellular growth.

In *E. coli* mainly two types of cold shock proteins are induced, class I and class II. The quantity of the class I cold shock proteins remain low at 37°C and gets induced at a high level during cold shock condition. Class I cold shock proteins are CspA, CspB, CspG, CspI, CsdA, PNP, RbfA, and NusA. Besides, the class II type of cold shock proteins remain at a certain extent at 37°C and gets induced at the moderate level during cold shock condition. Class II cold shock proteins are RecA, IF-2, H-NS, α subunit of DNA gyrase, Hsc66, dihydrolipoamide acetyltransferase, HscB, trigger factor, and pyruvate dehydrogenase (Rodriguez-Romo et al. 2005).

Among all the CSPs, the extensively studied major cold shock protein CspA functions as RNA chaperone. It prevents the formation of RNase-resistant stable secondary structures of the mRNA transcripts and thus facilitates its degradation at low temperature (Jiang *et al*., 1997). The interesting fact is that it has a long 5’UTR of 159 nucleotides (Tanabe *et al*., 1992). Giuliodori *et al*., 2010 showed that this 5’UTR acts as thermo sensor. Upon temperature downshift, the “cold-shock” 5’UTR structure rearrange itself such a way that the Shine Dargano sequence becomes available for the ribosome binding and thus more efficiently translated than the 37°C structure. Thus, this major cold shock protein gets induced while the translation of the other cellular protein gets hindered during cold shock condition.

CsdA (Jones *et al*, 1996) is one of the major cold shock proteins that acts as a DEAD-box RNA helicase (Turner *et al*., 2007). Other than CsdA, *E. coli* has four more DEAD-box proteins; DbpA (Diges and Uhlenbeck 2001; Fuller-Pace *et al*., 1993), RhlB (Py *et al*., 1996), RhlE (Bizebard and Ferlenghi 2004), and SrmB (Nishi *et al*., 1988). DEAD-box RNA helicase proteins are universal, found in prokaryotes, eukaryotes and archaea as well. Like all the DEAD-box RNA helicase proteins, these five *E. coli* DEAD-box RNA helicase proteins contain a core DEAD-box domain and have high sequence similarity with the eukaryotic eIF4A (Rogers *et al*., 2001; Du *et al*., 2002). Among these five DEAD-box RNA helicases in *E. coli*, CsdA is the only one which is induced at a high level during cold shock condition and acts as major cold shock protein. It utilizes energy by hydrolyzing ATP molecules to unwind RNA duplex at low temperature (Turner *et al*., 2007). CsdA protein is necessary for the cold adaption, and deletion of the *csdA* gene causes impaired growth at low temperatures in *E. coli* (Jones *et al*, 1996, Charollais *et al*., 2004). During cold-shock condition, CsdA protein is involved in the initiation of translation (Jones *et al*., 1996, Lu *et al*., 1999), degradation of mRNAs, (Khemici *et al*., 2004, Prud’homme-Généreux A, *et al*. 2004, Yamanaka and Inouye 2001) and the biogenesis of 50S ribosomal subunits in *E. coli* (Charollais *et al*., 2004, Peil *et al*., 2008). In the case of some pathogenic bacteria, like as, *Yersinia pseudotuberculosis* (Palonen *et al*., 2012) and *Salmonella typhimurium* (Rouf *et al*., 2011) CsdA protein is essential for the cold adaptation also. Not only in bacteria, but CsdA homolog plays an essential role during seed germination in *Arabidopsis thaliana* plant under cold stress condition (Liu *et al*., 2016). In addition, in an Antarctic archaeon, *Methanococcoidis burtonii*, CsdA protein homolog has been found to play an important role in growth and survival in its natural cold environment (Lim *et al*., 2000).

Despite the importance, the regulation of the *csdA* gene expression has not been studied in detail yet. About two decades ago, by primer extension method, Toone *et al*., 1991, observed two different lengths of *csdA* mRNA transcripts, one as major; 226 nucleotides and another as minor; 703 nucleotides. They also bioinformatically predicted a promoter region immediately upstream of 226 nucleotides long 5’UTR. On the basis of their study, later, Jiang *et al*., 1996 and Fang *et al*., 1998 studied the effect of the 226 bases long 5’UTR on its own expression. Fang *et al*., 1998, also found that the overproduction of 226 bases long 5’UTR causes derepression of its own gene along with the *cspA* gene. They assumed that this 226 bases may contain any repressor binding site.

In this study, we have focused on the identification of the promoter region experimentally and thus the actual length of the 5’UTR region of the *csdA* gene and its role in its gene regulation. Since the primer extension experiments by Toone *et al*., 1991, did not help conclusively in identifying the promoter region, we needed to find out the promoter of the *csdA* gene through other experimental means. For this purposes, we have constructed a promoter-less plasmid vector and found out that *csdA* promoter is present more than 800 bases upstream of the coding region of the *csdA* gene. Furthermore, we found that the remarkably long 5’UTR region of the *csdA* gene negatively regulates its own gene expression.

## Materials and Methods

### Green fluorescence intensity measurement

To measure the green fluorescence intensity, 1% of the overnight culture of *E. coli* was freshly inoculated in the LB media and the cells were grown till the mid-log phase (O.D. ~ 0.5-0.6) at 37 °C. When the cells reached the mid-log phase, the samples were pelleted down by centrifugation. The rest of the culture was transferred to 15°C and collected separately after 1hr of cold shock. The cells were re-suspended in 1ml of PBS with 2mM PMSF. The cells were then sonicated (sonication cycle specification; amplitude 100 %, 15 sec on and 20 sec off-cycle, for 3 minutes) followed by the pellet down by centrifugation at 13,000 rpm for 30 minutes at 4°C. The supernatant was collected and the total protein was quantified by using the standard Bradford method. 50 ug/ml of total protein in PBS (pH 7.4) was used to measure the green fluorescence intensity by fluorescent spectroscopy (excitation- 482nm, emission range- 490-600nm, slit-5).

### PCR amplification of *csdA* mRNA

The total cellular RNA was isolated using zymo research RNA isolation kit (#R2050) from the cells growing at the mid-log phase (O.D. ~ 0.5-0.6). The RNA samples were treated with DNaseI enzyme for 30 min at 37°C to remove any trace amount of contaminated genomic DNA (gDNA). DNaseI enzyme was inactivated at 65°C for 10 min in presence of EDTA. The quantity of the RNA was measured at OD260nm in spectrophotometer. 1ug of RNA was further used for the cDNA amplification using *csdA* mRNA coding region-specific reverse primer using thermo scientific cDNA synthesis kit (#K1621). The RNA samples were denatured at 65°C for 5min and cDNA was synthesized at 42°C for 1hr with the specifically designed reverse primer 5’AACAGAGGTAAAGAGAATGCTGCAGTTTTTCCGCTCCCC 3’ (3A, 3B, and 9A) and terminated at 70°C for 10 min. The synthesized cDNA was then amplified with specific and appropriate forward primers; 5’ AATGGAAAGGCTCAAGG 3’ (3A), 5’ CGCGGGATCGCATTATATTACGGCGGTCGTGACAAG 3’ (3B) 5’ GGCGTAAAAGTGAAGTCCTCG 3’ (9A). In each case, there was a control used in which cases the reverse transcriptase enzyme was not used to check if the sample was contaminated with gDNA or not.

### cDNA length study using denaturing 5% poly-acrylamide gel electrophoresis containing 8M urea

The reverse transcription reaction was performed from the total cellular RNA by using a reverse primer, which is complementary to the 194 bases downstream of the start codon of *csdA* gene. Before the reverse transcription reaction, contaminated DNA was removed with DNaseI treatment which was deactivated after the reaction. The specificity of this 39 bases long primer 5’AACAGAGGTAAAGAGAATGCTGCAGTTTTTCCGCTCCCC3’ was checked in the BLAST web server. After the cDNA synthesis, the reverse transcribed product was treated with the RNase A enzyme for 30 min at 37°C to remove the RNA samples. The amplified cDNA species were run on the denaturing 5% polyacrylamide gel electrophoresis containing 8M urea in 1X TAE buffer at 100V for 2hrs. Prior to the sample loading, all the cDNA samples were incubated at 95 °C for 5min and were chilled on ice immediately.

### EMSA experiment of *in vitro* RNA polymerase holoenzyme binding

All the DNA fragments were PCR amplified and ran on 1% agarose gel. The specific bands were purified using gel purification kit (Qiagen gel purification kit). All these DNA amplified products were *γ*-^32^P radiolabelled by using T4 polynucleotide kinase enzyme using [*γ*-32P] ATP as the substrate with the reaction condition of 37°C for 30 min followed by 65°C for 25min enzyme inactivation period. To remove the excess radioactivity, the samples were precipitated using double 100% ethanol and 1/10 volume of sodium acetate at pH 7.2. It was kept at −80°C for overnight. The samples were pellet down by centrifugation at 13,000 rpm for 30min at 4°C. The pellets were washed with 70% ice-chilled ethanol and the dried samples were re-suspended in nuclease-free water. The binding assay was performed by incubating these radiolabelled DNA fragments with RNA polymerase holoenzyme (NEB #MO551S) for 5min at 25 °C followed by loading in 4% polyacrylamide gel in 1X TBE buffer. The gel was run on 120V for 3hrs. Then it was dried in gel dryer (Bio-rad) and the image was taken in Typhoon TRIO + Variable Mode Imager.

### In *vivo csdA* promoter deletion

FRT-PGK-gb2-neo-FRT sequence amplification: The primers used for the *in vivo* deletion experiment were; forward primer- 5’ CTTTACAGCAGGAAGTGATTCTGGCACGTATGGAACAAATCCTTGCCAGTAATTAACCCTCACTAAAGGGCG 3’ and the reverse primer- 5’ CGCGATTCAAGTGCGCGTAGTTGTAAGTTGGATCAAGCTCAAGTACAGAATAATACGACTCACTATAGGGCTC 3’. In both of the cases, 22 nucleotides of 3’ends of these primers were complementary with the 5’ (forward primer) and 3’ end (reverse primer) of the FRT-PGK-gb2-neo-FRT sequence provided with K006 *E. coli* Gene Deletion Kit (Gene Bridges; Angrand et, al., 1999, Datsenko and Wanner 2000). Using these primers, the FRT-PGK-gb2-neo-FRT sequence was amplified to add 50 bases homology arms which corresponds to the sequences flanking the 200 bases *csdA* promoter region which was targeted for the replacement by FRT-PGK-gb2-neo-FRT sequence within the *E. coli* genome. The resulting fragment was gel purified.

Red/ET expression plasmid pRedET transformation: A single *E. coli* K-12 MG1655 colony was inoculated in 1.5 ml microfuge tube having 1 ml LB medium. A hole in the lid was punctured for the air and was incubated at 37°C overnight with 200 rpm shaking condition.From this 30 μl of the overnight culture was freshly inoculated into another fresh 1 ml LB medium and was grown for 2-3 hrs. at 37°C by shaking at 1000 rpm. The cells were centrifuged for 30 sec at 11,000 rpm at 4°C. The pellet was re-suspended with 1 ml chilled nuclease-free water. It was repeated twice and in the final, 30 μl was left in the tube with the pellet. 1 μl pRedET vector sample was mixed with the cell pellet and electroporated in 1 mm electroporation cuvette at 1350 V with a 5 ms pulse. The electroporated cells were resuspended in 1 ml LB medium without antibiotics and incubated at 30°C for 70 min by shaking at 1000 rpm. 100 μl cells were plated on LB agar plates containing ampicillin (50 μg/ml). The plates were incubated at 30°C for overnight.

Replacement of the *csdA* promoter region (200 bases) by the FRT-flanked PGK-gb2-neo cassette: A single colony containing pRedET plasmid from the ampicillin plate was inoculated in 1.0 ml LB media having 5ug/ml ampicillin and incubated at 30°C for overnight after puncturing a hole in its lid. 1.4 ml fresh LB media with ampicillin was inoculated with 30 μl of overnight culture and incubated at 30°C for 2 hrs by shaking at 1100 rpm. At OD600 0.3 of the culture, 10% L-arabinose was added to make the final concentration of 0.3%-0.4%. Then it was incubated at 37°C, shaking for 45 min to 1 h to induce the expression of the Red/ET recombination proteins. The cells were centrifuged for 30 sec at 11,000 rpm at 4°C. The pellet was re-suspended with 1 ml chilled nuclease-free water. These processes were repeated and in final, 30 μl was left in the tube with the pellet. 1-2 μl (200 ng) of amplified (purified) linear FRT-PGK-gb2-neo-FRT fragment with homology arms was added and electroporated as discussed above. 1 ml of LB medium without antibiotics was added to the cuvette and mixed carefully by pipetting. The culture was incubated at 37°C with shaking for 3 hours. The culture was then pelleted down with the remaining medium and a loop was streaked onto LB agar plates containing kanamycin (15 μg/ml) but no ampicillin. The plate was incubated at 37°C overnight at which the Red/ET recombination protein expression plasmid (pRedET) disappeared.

Confirmation of successfull replacement of *csdA* promoter with FRT-PGK-gb2-neo-FRT fragment within the *E. coli* genome: One of the colonies from the kanamycin plate was picked and the genomic DNA was isolated from the overnight grown culture. We did sequencing for the confirmation of the correct insertion inside the *E. coli* chromosome by using a forward primer (5’ TATCAGGACATAGCGTTGGCTACC 3’), which was complementary to a region located on the FRT-PGK-gb2-neo-FRT cassette, upstream of the 50 bases homology region at the 3’ end of the 200 bases *csdA* promoter fragment. A reverse primer was used (5’ CGAGACTAGTGAGACGTGCTAC 3’) which was also complementary to a region located on the FRT-PGK-gb2-neo-FRT cassette but downstream of the 50 bases homology region at the 5’ end of the 200 bases *csdA* promoter fragment. Both these sequence confirmation results have been provided in supplementary figure 1.

### Western blot experiment using CsdA antibody

We have purified CsdA protein to raise polyclonal antibody against it. We had hyper-immunized a New Zealand White rabbit with 90ug of purified CsdA protein mixed with Complete Freund’s adjuvant (CFA). Half of the initial protein amount was further immunized mixed with the Incomplete Freund’s adjuvant (IFA) for next two weeks. The blood sample was collected after 7 days of the third immunization. Then we performed Western blot analysis to confirm the antibody production against the CsdA protein and to standardize the amount of the rabbit serum (primary antibody (1:5000)). For the western blot experiment, equal amount of the samples were loaded on the SDS-PAGE gel and after the run the protein bands were transferred to membrane followed by visualized using Ponceau S staining (supplementary figure 2).

## Results

### 1.1 Modification in commercially available pTurboGFP-B vector

We have modified the commercially available pTurboGFP–B vector (catalog no. # FP513) (Fig. 1A). This vector originally contains T5 promoter/lac operator element to express the green fluorescence protein (GFP). We have performed two step modifications in this vector to use it as promoter less vector for the purpose of the identification of *csdA* promoter region. In the first step, to generate EcoRI restriction site, we have inserted single nucleotide “C” in the 93^rd^ position immediately downstream of T5 promoter/lac operator element by the process of site-directed PCR based mutagenesis. Then, in the second step, by using XhoI (already present in the original vector) and newly introduced EcoRI restriction site, we have replaced T5 promoter/lac operator region with a DNA fragment which contains multiple cloning site (MCS) region containing XhoI, BglII, EcoRV, XmaI, SmaI, KpnI, Eco53kI, SacI, HpaI, NotI, SacII, SalI, and EcoRI from the 5’ to 3’ direction (Fig. 1B). Thus, the resultant modified pTurboGFP–B vector lacks the promoter region but have a MCS region instead. Afterward, we have inserted the promoter region of the major cold shock gene, *cspA* within the MCS region of this modified pTurboGFP – B vector using BglII and SalI restriction enzymes. We have measured the green fluorescence intensity of these two constructs. We have not found any green fluorescence intensity in modified pTurboGFP–B vector with MCS region, in contrast, we have found green fluorescence intensity in modified pTurboGFP–B vector with *cspA* promoter region (Fig. 1C).

**Fig. 1:**
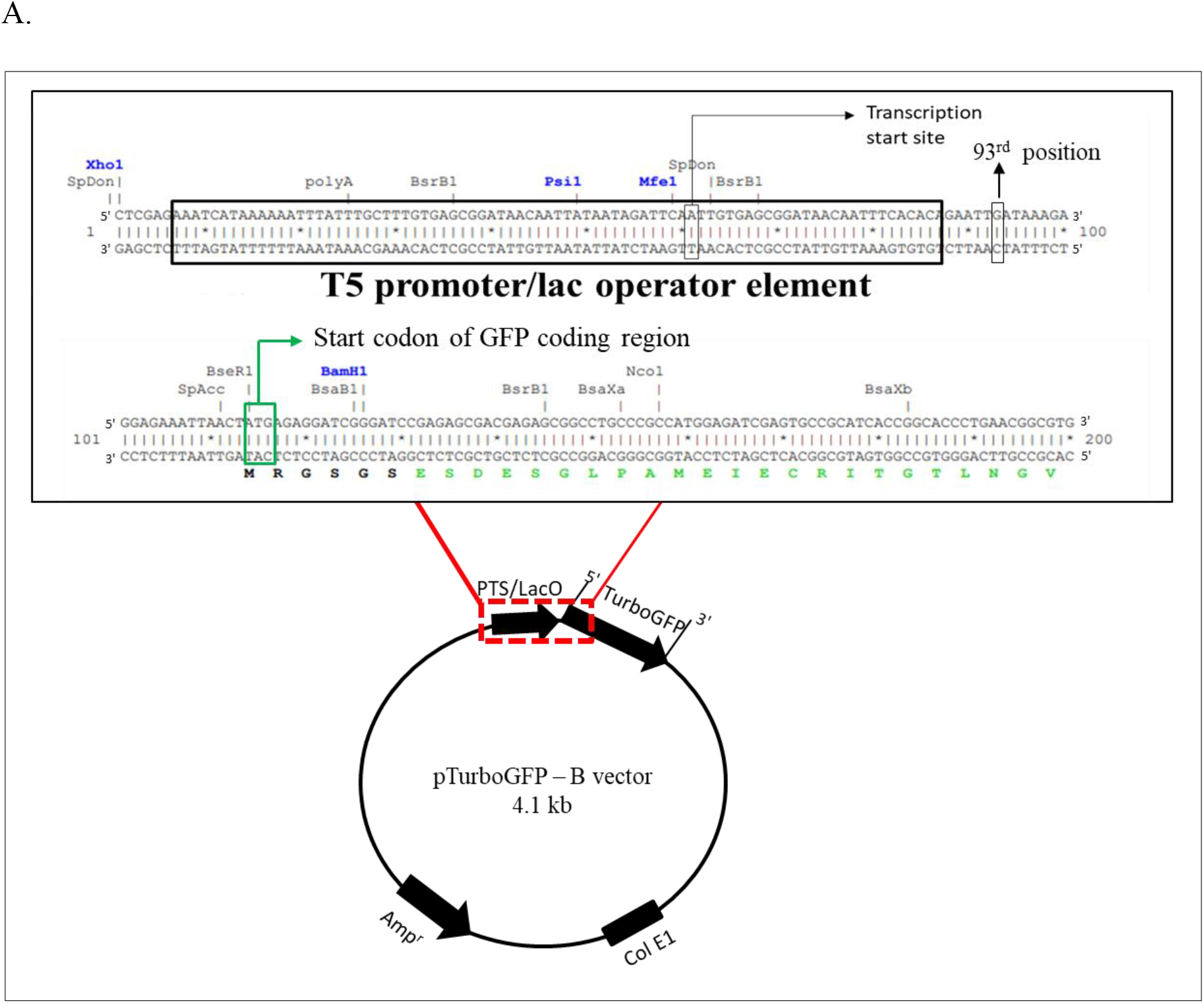
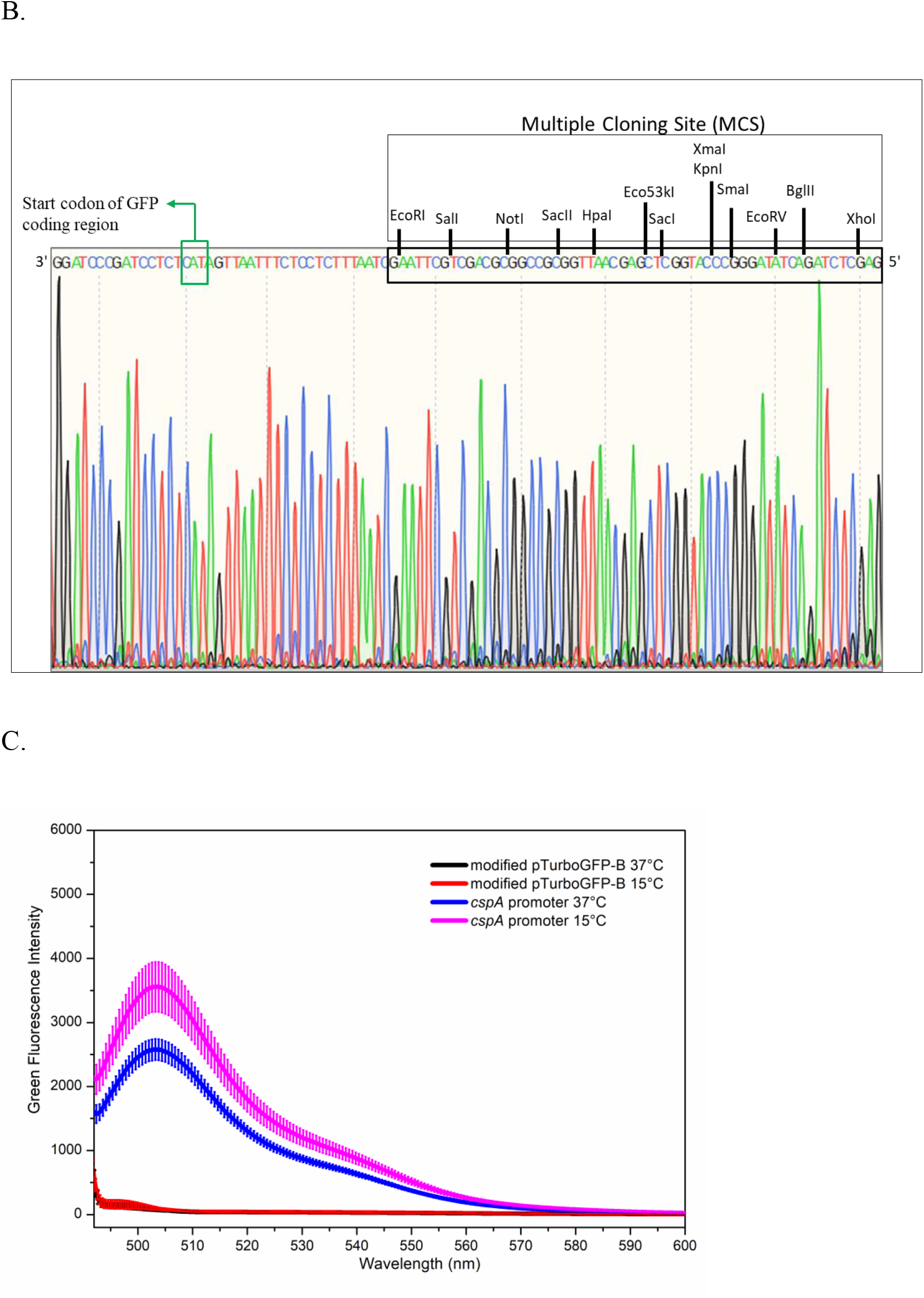
Construction of modified pTurboGFP–B vector by the replacement of T5 promoter/lac operator element of original pTurboGFP–B vector with MCS region. (A) Graphical representation of original pTurboGFP–B vector. The sequences of the T5 promoter/lac operator element (PTS/LacO) along with the upstream and downstream regions of the original pTurboGFP–B vector is shown in detail in red dotted box. T5 promoter/lac operator element is shown in black box. The transcription start site (black) and the translation start codon (green) of the GFP coding region are marked by the arrow. The 93^rd^ position at which the “C” nucleotide is inserted has been marked. (B) Sequence confirmation of modified pTurboGFP-B vector containing MCS region instead of T5 promoter/lac operator element. MCS region is shown in the black box. The restriction sites present within the MCS region are shown sequentially. The translation start codon of the GFP coding region is also marked. (C) Green fluorescence intensity measurement of modified pTurboGFP-B vector with MCS and *cspA* promoter region. Modified pTurboGFP–B vector with *cspA* promoter region is showing green fluorescence intensity both at 37°C and 15°C also in contrast, modified pTurboGFP-B vector with MCS region is showing no green fluorescence intensity.

### 1.2 No promoter region found immediate upstream of the coding region of *csdA* gene

We have amplified 194 base fragment encompassing the Toone *et al*. predicted −35 region and −10 region immediately upstream of 226 bases 5’UTR. By the same procedure, like as *cspA* promoter cloning, we have inserted this 194 base fragment using BglII and SalI restriction enzymes in the MCS region of this modified pTurboGFP-B vector. With respect to the positive control, we have not found any green fluorescence intensity in this 194 base fragment, (Fig. 2A). Further, in search of *csdA* promoter, we have cloned 103 bases fragment immediate upstream of *csdA* coding region followed up to 700 bases by adding approximately 100 upstream bases each time (103 bases (Cod 103bpU), 208 bases (Cod 208bpU), 320 bases (Cod 320bpU), 414 bases (Cod 414bpU), 540 bases (Cod 540bpU), and 700 bases (Cod 700bpU)) in modified pTurboGFP – B vector using the same BglII and SalI restriction sites. Fig. 2B showed that there is no green fluorescence activity up to 700 bases region immediately upstream of *csdA* coding region with respect to the promoter of *cspA*, positive control.

**Fig. 2.**
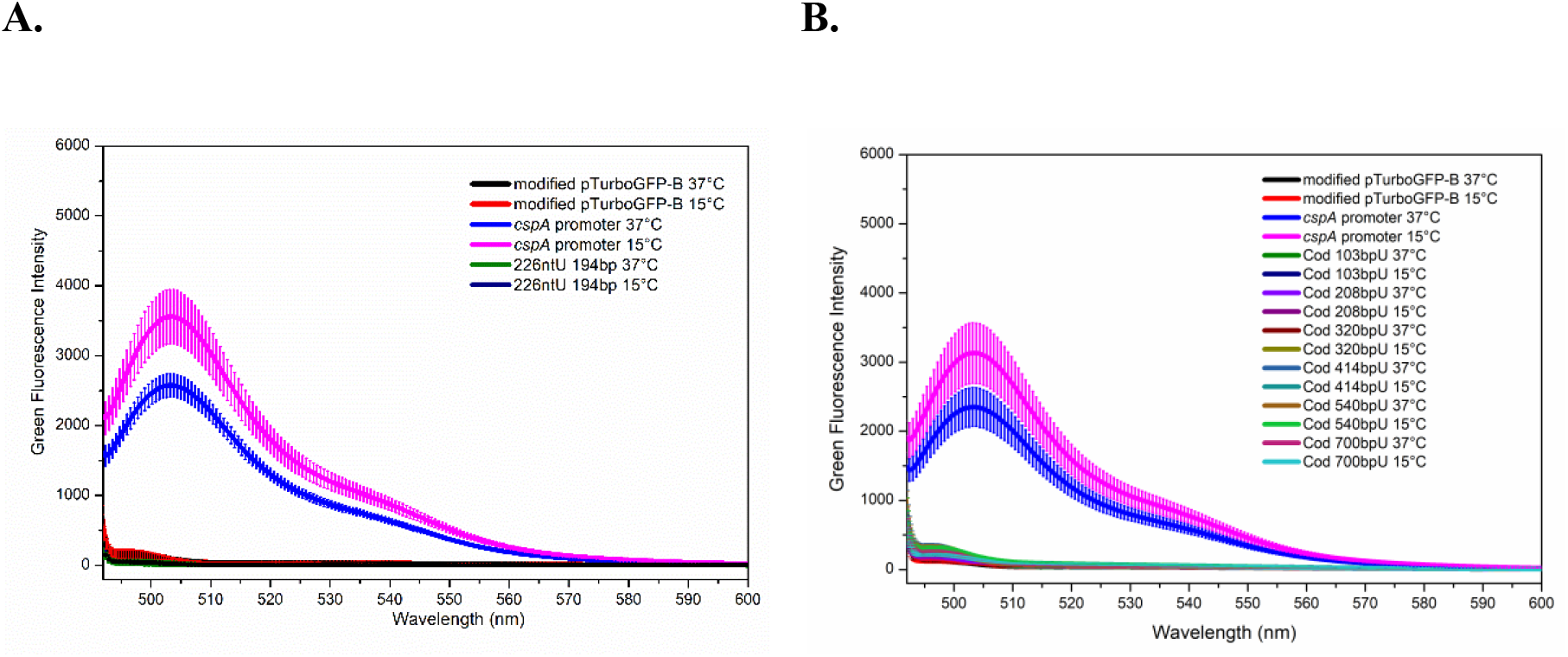
Emission spectra scanning of green fluorescence intensity. (A) No green fluorescence is seen in the 194 base fragment immediate upstream of 226 nucleotides long 5’UTR with respect to the positive control, *cspA* promoter containing modified pTurboGFP-B vector. (B) 103 bases, 208 bases, 320 bases, 414 bases, 540 bases, and 700 bases fragments immediate upstream of *csdA* coding region are also showing no green fluorescence intensity with respect to the positive control.

### 1.3 *In vivo* evidence of remarkably long 5’UTR

Toone *et al*. 1991., did not further mention the 703 bases long 5’UTR as it was found as minor species. Further, several studies followed the finding of 226 bases 5’UTR (major species) as they bioinformatically predicted a promoter region immediate upstream of this 5’UTR region. However, as shown in Fig. 2A, our *in vitro* studies have found no promoter activity in this predicted promoter region. We further tested if the *csdA* mRNA transcript is longer than 226 bases or not by *in vivo* approaches. We have used a reverse primer complementary to the *csdA* coding region (194 bases downstream of the start codon) to reverse transcribe the *csdA* mRNA specifically, among the total cellular RNA of *E. coli*. Fig. 3A is showing the amplified product of 614 bases fragment using the forward primer complementary to the 194 bases upstream of the 226 bases long 5’UTR from genomic DNA (positive control), total cellular RNA (DNase I treated to check the gDNA contamination), and cDNA (reverse transcribed using above-mentioned *csdA* coding region-specific reverse primer). Further, to find out if the length of mRNA transcript is approximately 703 bases long or not, we isolated total cellular RNA and PCR-amplified *csdA* mRNA specifically using the same reverse primer along with a forward primer complementary to the 44 bases downstream from the start point of 703nt transcript. Our results showing the amplified product of 853 bases from the wild-type genomic DNA (positive control) and cDNA but not from the total cellular RNA (Fig. 3B), have established the fact that *csdA* mRNA is atleast about 700 bases long.

**Fig. 3:**
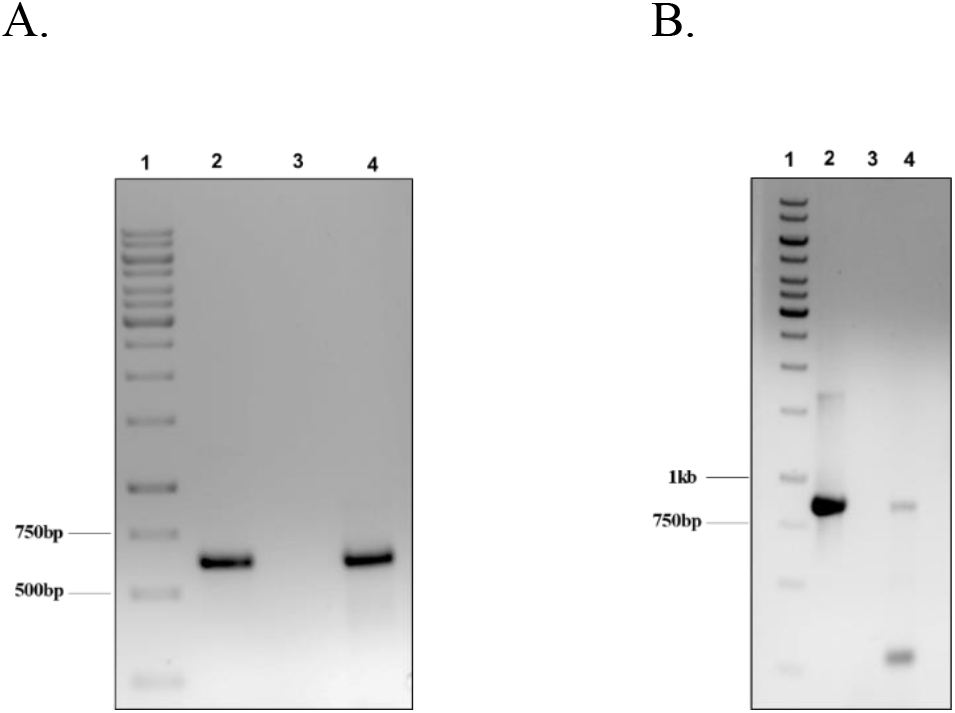
Ethidium bromide–stained 1% agarose gel of PCR-amplified products. Lane 1 represents the DNA ladder. Lane-2: wild-type genomic DNA. Lane-3: RNA template. Lane-4: cDNA. (A) Length of the amplified product is 614 bases. (B) Length of the amplified product is 853 bases.

### 1.4 *In vitro* evidence of *csdA* promoter 700 bases upstream of its coding region

In search of *csdA* promoter upstream of the 703 bases 5’UTR, we have amplified 53 bases immediately upstream of 703 bases long 5’UTR using BglII and SalI restriction enzymes in the modified pTurboGFP – B vector. We have not found any green fluorescence intensity (Fig. 4A) using this fragment. Similarly, we have amplified 100 bases, 150 bases, and 200 bases upstream of the −703 nt fragments in the modified pTurboGFP – B vector. Fig. 4A is showing that rather than that of 100 base and 150 base fragments, 200 base fragment has the promoter activity both at 37°C and 15°C. No green fluorescence activity from the fragments of 50 bases, 100 bases, and 150 bases indicates that 5’UTR is longer than 703 bases. Besides, we have performed EMSA experiment to test if in the *in vitro* condition, the RNA polymerase holoenzyme (includes σ70 factor) can bind to this *csdA* promoter region (200 bases) or not. In this experiment, along with the 50 bases, 100 bases, 150 bases, and 200 bases fragments, we have used *cspA* promoter region (133 bases) as positive control and *csdA* coding region fragment as the negative control. We have found that at the 25°C reaction temperature for 5 minute incubation time period, the RNA polymerase holoenzyme is not binding to the negative control but to the positive control and the experimental cases (Fig. 4B). It is indicating that RNA polymerase holoenzyme has high affinity to this *in vitro* identified promoter region.

**Fig. 4:**
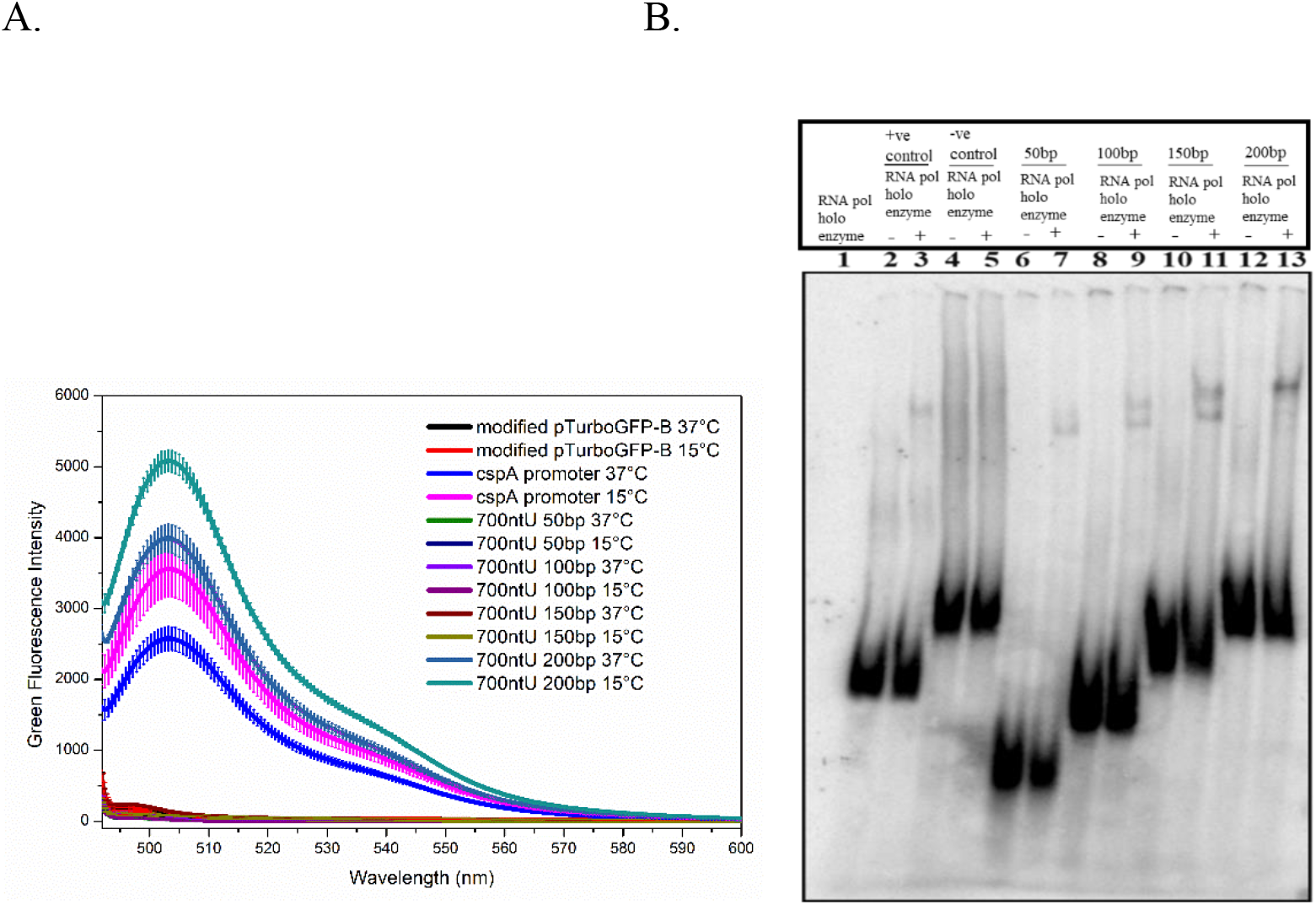
Green fluorescence intensity of the fragments 700 bases upstream of the coding region of *csdA* gene. (A) Like as modified pTurboGFP-B vector (negative control), 50 base, 100 base and150 base fragments 700 bases fragments upstream of *csdA* coding region are not showing any green fluorescence intensity. On the other hand, 200 base fragment is showing green fluorescence intensity both at 37°C and 15°C. (B) EMSA experiment of RNA polymerase holoenzyme binding to the promoter fragments on 4% polyacrylamide gel. Lane 1; *E. coli* RNA polymerase holoenzyme [non-radio [γ-^32^P] labelled]. Lane 2 and 3 - *cspA* promoter, Lane 4 and 5 - *csdA* coding region, Lane 6 and 7 - 50 base fragment, Lane 8 and 9 - 100 base fragment, Lane 10 and 11 - 150 base fragment, Lane 12 and 13 - 200 base fragment, without and with RNA polymerase holoenzyme respectively. All these fragments are [*γ*-32P] radiolabeled.

### 1.5 *In vivo* deletion of *csdA* promoter region from *E. coli* genome

Based on these *in vitro* studies, to approve the promoter activity of this 200 base fragment *in vivo* context as well, we have deleted this fragment from the *E. coli* genome. For this purpose, we have used the Red/ET recombination tool developed by gene bridges (Cat. No. K006). The detailed protocol has been described in the material and method section. We have confirmed the accuracy of deletion by the sequencing results provided in supplementary figure 1. We have used in house built CsdA antibody to detect the cellular CsdA protein in that 200 bases deleted mutant strain with respect to the wild type strain. We have observed that the deletion causes decrease in the CsdA protein level in the mutant type both at 37°C and 15 °C with respect to the wild type (Fig. 5A and 5B). During the cold shock condition (15°C), we have found that there is approximately five times reduction of CsdA protein level in the mutant strain with respect to the wild type (Fig. 5B).

**Fig. 5.**
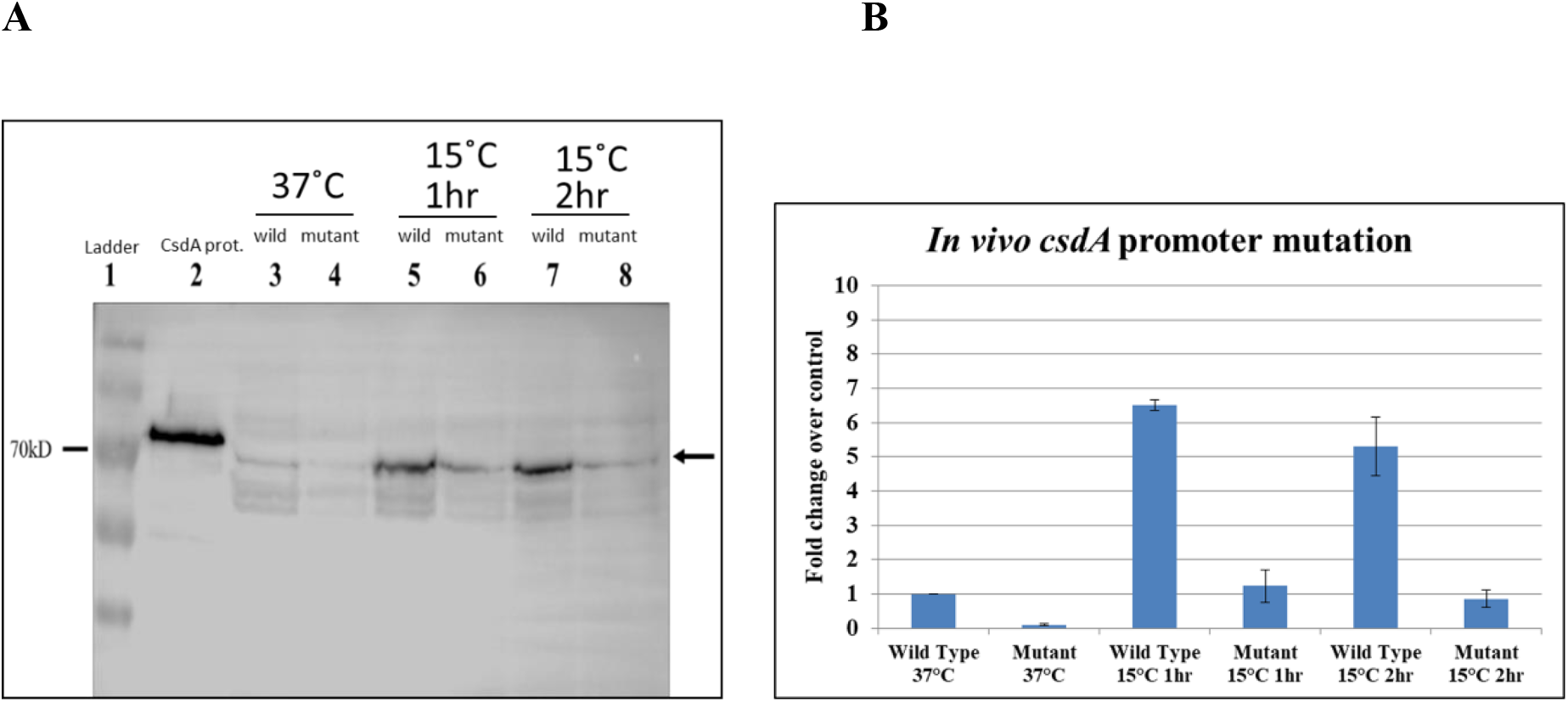
Cellular CsdA protein level detection with CsdA antibody in wild type and *csdA* promoter deleted mutant *E. coli* strain. (A) Western blotting with CsdA antibody. Lane 1 – protein ladder, Lane 2 – purified His-tag CsdA protein, Lane 3, 5 and 7 - CsdA protein of wild type *E. coli* at 37°C, 1hr at 15°C, and 2hr at 15°C respectively. Lane – 4, 6, and 8 CsdA protein of mutant type *E. coli* at 37°C, 1hr at 15°C, and 2hr at 15°C respectively. The black arrow is indicating CsdA antibody detected the CsdA protein band. (B) Bar diagram of CsdA protein expression fold change over control (wild type 37°C) and 200 base fragment deleted *E. coli* str.K12 substr.MG1655 at 37°C and 15°C along with standard deviation values.

### 1.6 Identification of −10 region and −35 region

To find out the exact location of the −10 region and −35 region within the 200 bases *csdA* promoter region, we have further amplified Δ50 fragment (50 bases) and Δ100 fragment (100 bases) which reside in between the 150 bases fragment to 200 bases fragment and 100 bases fragment to 200 bases fragment respectively. Δ100 fragment has shown measurable amount of green fluorescence intensity both at 37°C and 15°C whereas Δ50 fragment did not show any green fluorescence intensity (Fig. 6A). To determine the sequence of the −10 region and - 35 region present within the Δ100 fragment, we have aligned this fragment with the known promoters of other major cold-shock genes *cspA*, *cspB*, *cspG*, and *cspI* of *E. coli* (Fig. 6B).

**Fig. 6.**
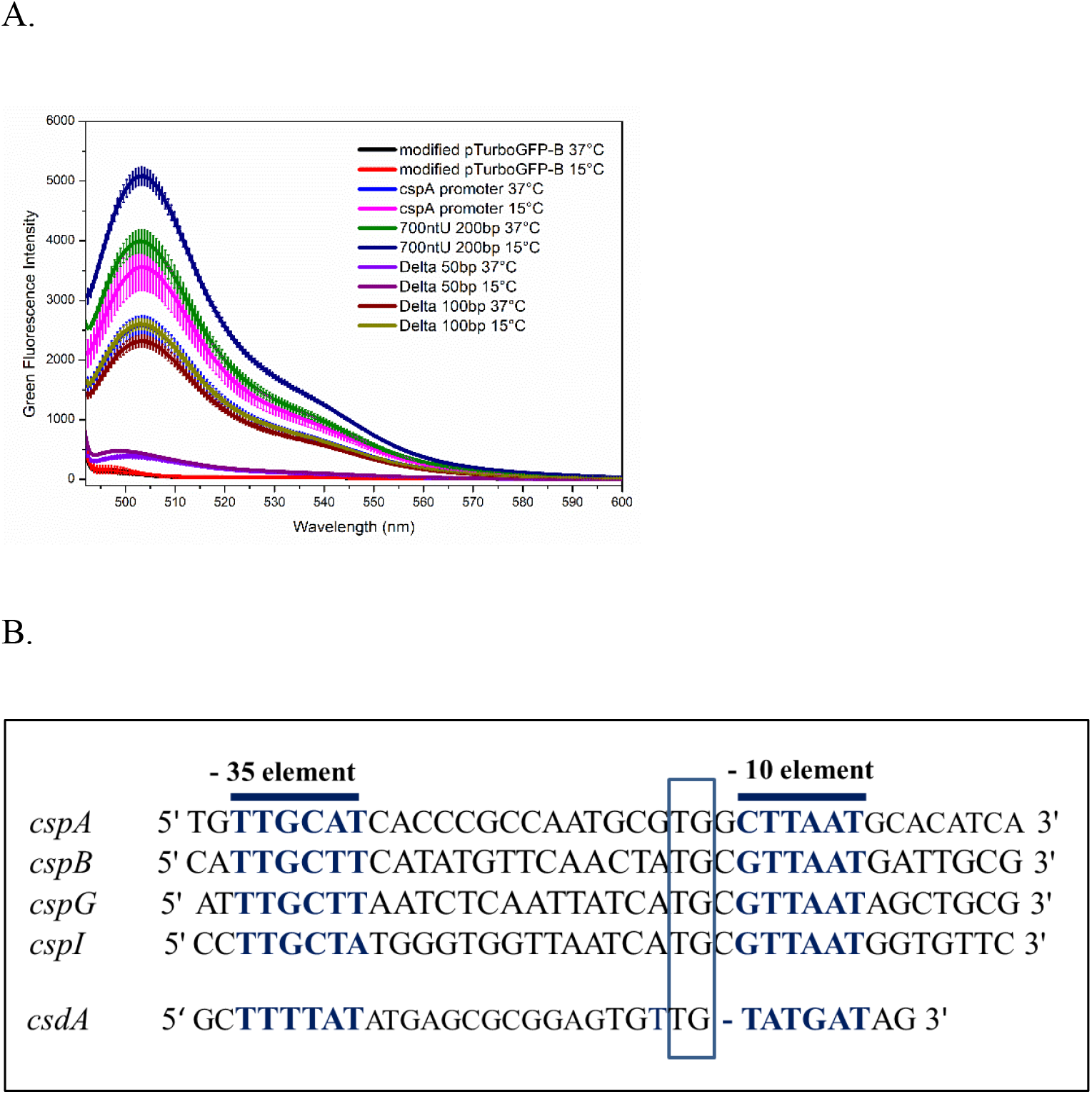
Identification of −35 element and −10 element: (A) Green fluorescence intensity of the Δ50 fragment and Δ100 fragments. Δ50 does not show green fluorescence but Δ100 showed green fluorescence. (B) Alignment of *csdA* promoter region with the promoter elements of other major cold-shock genes. −35 element and −10 element of *cspA*, *cspB*, *cspG*, and *cspI* are marked in bold and blue color. TGn motif is shown in the blue box.

Additionally, in case of major cold shock protein genes, *cspA* and its homologs *cspB*, *cspG*, and *cspI* (Ermolenko *et al*., 2002; Phadtare, S. 2004.) a TGn motif is present in their promoter region along with the −35 element and −10 element (Phadtare S and Severinov K, 2005). This TGn motif specifically interacts with an additional conserved region within σ^70^, called 3.0 (Campbell *et al*., 2002; Sanderson *et al*., 2003). Thus, in this study we have identified the - 35 sequence and the - 10 sequence, and the TGn motif also within Δ100 fragment, comparing it with the promoter of those cold-shock inducible genes (Figure. 5B). Recently, in 2020, the findings of −10 region and −35 region sequences of the *csdA* promoter by Ojha. S and Jain. C, are also supporting our observation. In addition, their 5’RACE experiment has found that the transcription of *csdA* gene starts 838 nucleotides upstream of the *csdA* start codon supporting our experimental results.

### 1.7 Role of the 5’UTR of *csdA* gene in its own regulation

The Δ100 promoter fragment itself contains 38 nucleotides of the 5’UTR region. The addition of another 100 nucleotides of the 5’UTR region to this Δ100 promoter fragment i.e., 200 base fragment (Fig. 4A) has shown increase in the green fluorescence intensity. But, further addition of another upstream 100 nucleotides causes decrease of green fluorescence intensity with comparison to the Δ100 fragment (Fig. 7). Furthermore, the sequential addition of upstream 100 nucleotides causes complete repression of green fluorescence intensity (Fig. 7). Furthermore, we found that the whole 5’UTR region (838 nucleotides) completely represses the expression of green fluorescence intensity (Fig. 7).

**Fig. 7.**
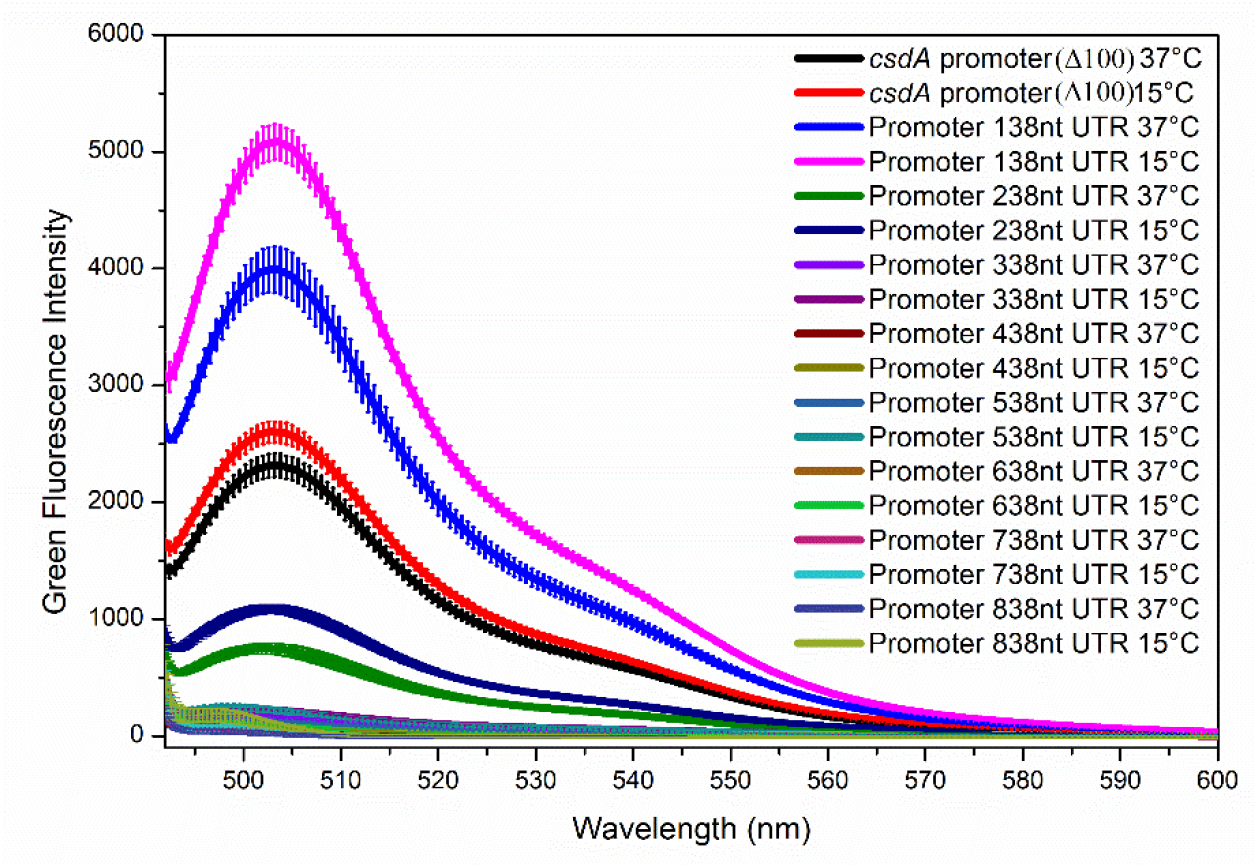
Green fluorescence intensity of the 5’UTR regions. 138 nucleotides, 238 nucleotides, 338 nucleotides, 438 nucleotides, 538 nucleotides, 638 nucleotides, 738 nucleotides, and the whole 838 nucleotides 5’UTR region including the first 5 codons of *csdA* coding region along with *csdA* promoter (Δ100 fragments) are shown with the standard deviation values.

### 1.8 *yrbN* gene inside the *csdA* gene 5’UTR region

In 2008, Hemm *et al*., observed a short ORF region which encodes 26 amino acids containing peptide while they were studying genes encoding small proteins (16–50 amino acids) in the intergenic regions of the *E. coli* genome by using sequence conservation models. This gene is named as *yrbN* gene. The function of this gene is still not known. In this study we have found that the nucleotide sequence of the coding region as well as the amino acid sequence of *yrbN* gene of bacteria belonging to the class of gammaproteobacteria, are conserved (Fig. 8A and B). Furthermore, the ORF region of *yrbN* and *csdA* overlaps with each other (Fig. 8C) but both of these ORFs are not present in the same frame (Fig 8C). We have previously described in figure 6 that the fragment containing the promoter region along with 738 nucleotide 5’UTR and another fragment containing 838 nucleotides 5’UTR fragments along with 5 codons of *csdA* coding region; both of these showed no green fluorescence. As the complete *yrbN* gene resides within the region in between 738 nucleotide 5’UTR fragment and 838 nucleotide 5’UTR fragment along with 5 codons of *csdA* coding region (Fig. 8D), we can conclude that *yrbN* gene does affect the repression activity of the whole 5’UTR region.

**Fig. 8:**
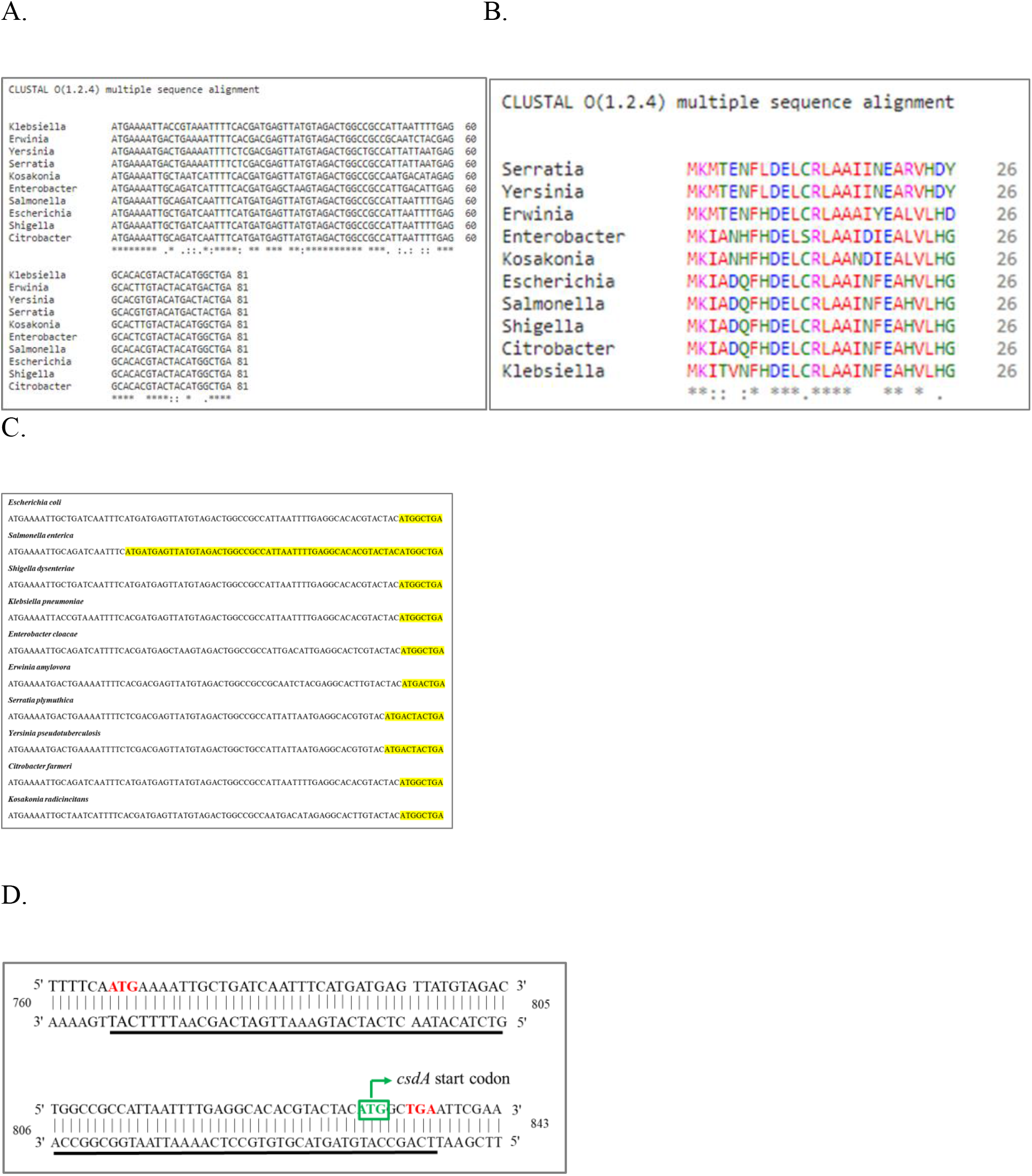
Study of *yrbN* gene in *E. coli* and gammaproteobateria. (A) Alignments of ORFs of the *yrbN* gene of the bacteria belong to gammaproteobacteria. (B) Alignments of amino acid sequences *yrbN* gene of those bacteria. All these sequences are taken from the NCBI database and Uniprot database and have been aligned by using ClustalW2. ‘*’ indicates identical residues and ‘:’ and ‘.’ indicate conserved and semiconservative substitutions respectively. (C) Overlapping region of *csdA* and *yrbN* gene coding region. The overlapping nucleotides have been highlighted with yellow color. In all these cases, the ORF of the *yrbN* gene resides within the 5’UTR of the *csdA* gene. (D) The start codon and stop codon of *yrbN* gene are shown in red colour. *yrbN* gene ORF region is marked in black line. Start codon of *csdA* gene is shown in green box. This 5’UTR region is numbered from the transcription start site (TSS) as +1.

### 1.9 Evidence of *csdA* expression from *nlpI* gene and *pnp* gene

Our *in vivo* deletion experiment of *csdA* promoter region (Fig. 5A) from the *E. coli* genome revealed that the deletion could not completely abolish the expression of *csdA* gene. This may indicate the fact that *csdA* gene expression may also happen from the promoter region of the immediate upstream *nlpI* gene. Thus, we have used a forward primer which is 89 bases upstream of the Δ100 *csdA* promoter region, and a reverse primer 194 bases downstream of *csdA* coding region to amplify the *csdA* specific mRNA. We have found a 1186 bases long amplified product (Fig. 9A). Furthermore, we have reverse transcribed the *csdA* specific mRNA using that same reverse primer 194 bases downstream of the start codon of *csdA* gene and run the cDNA on the denaturing 5% polyacrylamide gel electrophoresis containing 8M urea. We have found an approximately 1000 nucleotide long cDNA product both at 37°C and 15°C. Here, we have used the reverse primer 194 nucleotides downstream of the *csdA* coding region. Thus, this cDNA product is approximately 806 nucleotides long (i.e., it can be 838 bases long 5’UTR transcribes from *csdA* promoter that we have identified). We have also found two other cDNA products larger than 1000 nucleotides (Fig. 9B). One of these products (second largest) was present at both the 37°C and 15°C samples. As the start codon of the *nlpI* gene is 1064 nucleotides upstream of the start codon of the *csdA* coding region (promoter and the transcription start site of *nlpI* gene are still not known), the approximate length of this transcript (Fig. 9B) indicating that the *csdA* gene may also be transcribed along with the *nlpI* gene, the immediate upstream gene of the *csdA* gene. There was another cDNA product (largest) only present at 15°C sample (Fig. 9B). As the transcription start site of the *pnp* gene is 3466 nucleotides upstream of the start codon of *csdA* coding region, the *csdA* mRNA may also transcribed from the promoter of the *pnp* gene during cold-shock condition.

**Fig. 9.**
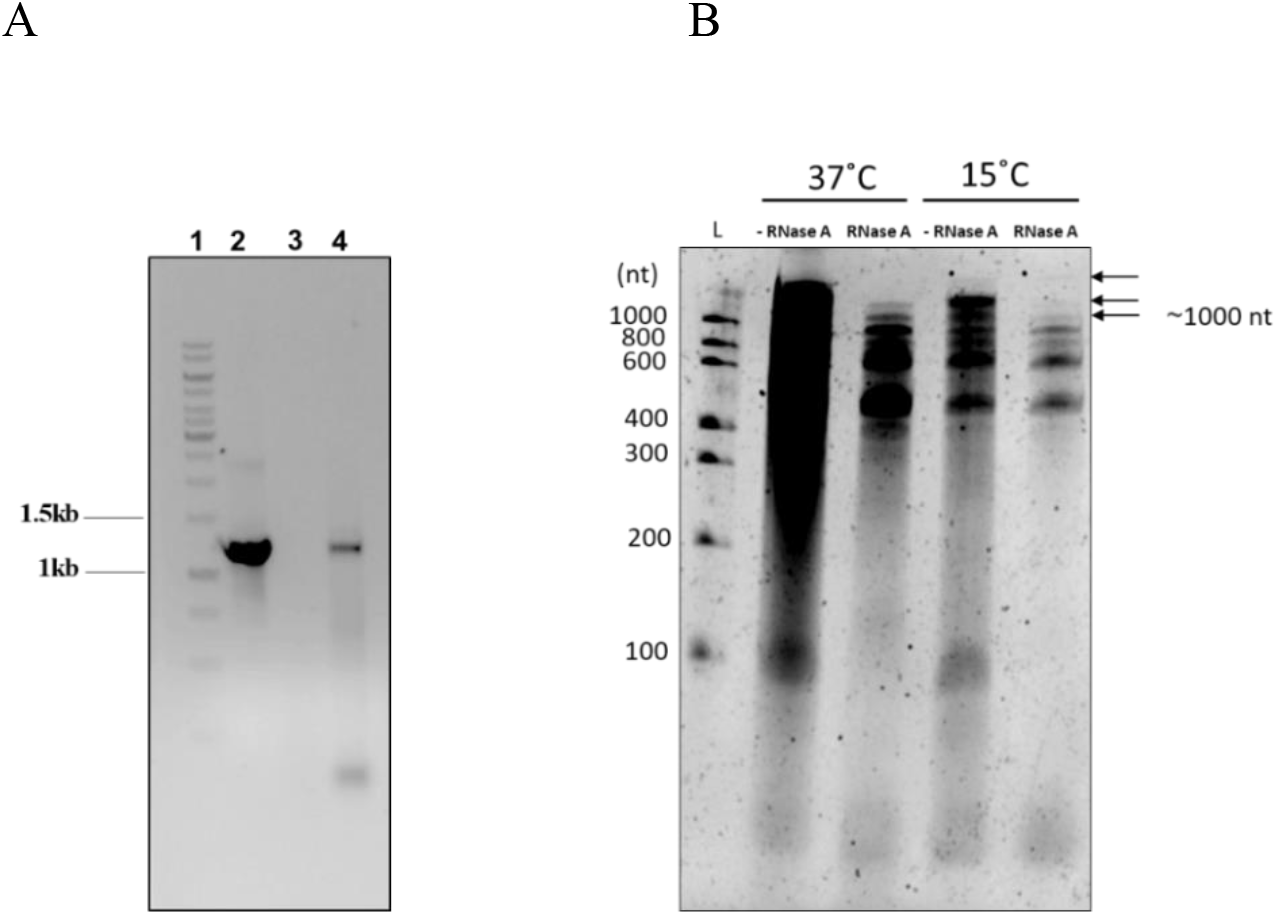
PCR amplified products and reverse transcribed products of the *csdA* mRNA. (A) PCR amplified products of the *csdA* mRNA. Lane 1 is the DNA ladder. Lane 2 and 4 indicate the fragment amplified from the gDNA and cDNA of *E. coli*. Lane 3 is the amplified product of the mRNA. The amplified product length is 1184 bases. (B) Urea-polyacrylamide gel image of the reverse transcribed product of the *csdA* mRNA. Lane 1 is the RNA ladder. Lane 2 and lane 4 are the cDNA samples not treated with the RNase A enzyme. Lane 3 and lane 5 are the cDNA samples treated with the RNase A enzyme.

## Discussion

As Toone *et al*., 1991, observed 226 bases long 5’UTR as major species and bioinformatically found a promoter region immediate upstream of this 5’UTR region, many studies further followed their observation. Though they also found 703 bases long 5’UTR as minor species, they did not further discuss it. But our study has proven that there is no promoter present immediate upstream of 226 bases long 5’UTR. Moreover, the 5’UTR is not 703 bases long but contains 838 bases (Ojha and Jain, 2020). In support of our study, we found an article published by Thomason *et al*. in 2015. They also found a transcription start site at the 838 bases upstream of the *csdA* gene coding region. Thus, the observations of the 226 bases major *csdA* mRNA species and 703 *csdA* mRNA minor species results of the primer extension study by Toone *et al*., may be due to the non-specific RNA breakage.

Now, if we focus on the *csdA* gene; we can find that the CsdA homolog is present in bacteria, eukaryotes, and archaea as well (Palonen *et al*., 2012, Rouf *et al*., 2011, Liu *et al*., 2016, Lim *et al*., 2000). Further studies revealed that 5’UTR of the *csdA* gene is remarkably long not only in bacteria but in archaea also. For example, in archaea, *Thermococcus kodakarensis*, expresses cold stress-inducible DEAD-box RNA helicase (Tk-deaD), which has a long 5’UTR of 158 bases (Nagaoka *et al*., 2013). Another Antarctic methanogen, *Methanococcoides burtonii* contains *dead* gene, which has 113 bases long 5’UTR (Lim *et al*., 2000). However, the role of these long 5’UTR regions in the regulation on its own gene expression has not been studied in details.

Not only *csdA* gene, but other major cold-shock genes *cspA*, *cspB*, *cspG*, and *cspI* in *E. coli*, also contain long 5’UTR (159 bases, 161 bases, 156 bases, and 145 bases respectively) (Tanabe *et al*., 1992; Etchegaray *et al*., 1996; Nakashima *et al*., 1996; Wang *et al*., 1999 respectively). It seems that it may be a common feature of the major cold-shock genes to contain a long 5’UTR. Surprisingly, we have found that the 5’UTR of the *csdA* gene is 838 bases long, which is remarkably long with respect to the length of the 5’UTR of the previously studied cold shock genes in *E. coli*. But when we have expressed this whole 5’UTR along with its promoter plus its five codons, we have found out that this whole 5’UTR actually represses its own gene expression. Then, the question rises if the 5’UTR does not contribute to the induction during cold temperature, then how it gets induced? Thus, we think that the reason for the cold induction may be due to the expression of the *csdA* gene from the promoter of the *pnp* gene during low temperature (Fig. 9B).

One of the common features of prokaryotes is the operon system. In this system, multiple genes are encoded by the single promoter region. As we have not found any green fluorescence intensity from any fragment downstream of the *csdA* promoter region, it is obvious that *yrbN* gene and *csdA* gene reside in an operon. The length of this *yrbN* gene ORF is only 81 bases and it encodes a 26 amino acid short peptide. It seems this short peptide may act as a leader peptide. Moreover, we found that the 5’UTR region of the *csdA* gene without the *yrbN* gene ORF and the whole 5’UTR region of the *csdA* gene along with the *yrbN* gene ORF; both repress the *csdA* gene expression. It means *yrbN* gene may also represses the expression of *csdA* gene or it does not affect the repression activity of this 5’UTR region in this *csdA* gene expression. Further study is needed to explore the function of the protein encoded by this *yrbN* gene and the role of this gene in the regulation of the *csdA* gene expression.

In this study, we have observed multifarious types of regulation of the *csdA* gene expression (Fig. 10). Firstly, it has a remarkably long 5’UTR, which is uncommon in prokaryotes. Secondly, there is a short ORF region of the *yrbN* gene that resides within this long 5’UTR. Thirdly, *csdA* gene expression also occurs from the upstream *nlpI* (both at 37°C and 15°C) and *pnp* gene (only at 15°C) as well. All these phenomena of the multifarious regulation system may be related to the multiple functions of this CsdA protein. In 2004, Bizebard *et al*., observed that CsdA protein catalyzes ATP hydrolysis in the presence of different RNA substrates. In 1996, Jones *et al*., found that when the cells were grown at 37°C, CsdA is present in small quantities but induced significantly upon cold shock condition. As this protein consumes cellular energy in the form of ATP during low temperature (i.e., survival crisis) the cell may need confined regulation systems (especially the repression of the *csdA* gene by its 838 bases remarkably long 5’UTR) for the proper utilization of the energy during this crisis.

**Fig. 10.**
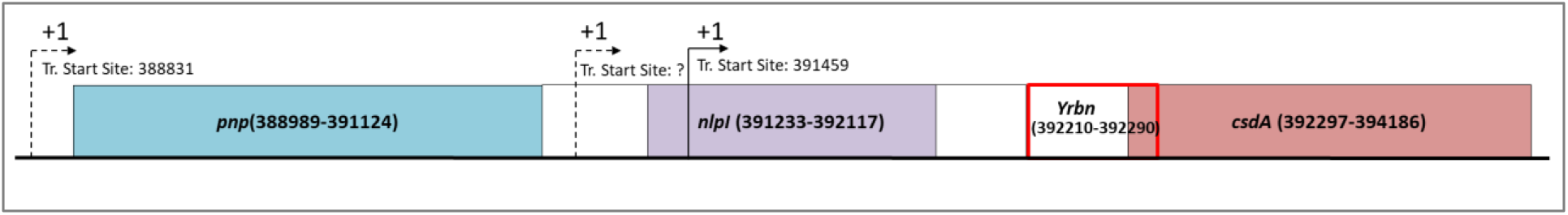
Graphical representation of the summary of this study. Transcription start sites of *csdA* gene are marked with arrow. The dotted arrow means it is not established yet. It is numbered as *Escherichia coli* str. K-12 substr. MG1655 with the ref: accession no. CP009685.

## Data Availability

All data supporting the findings in this study are provided in the main text and supplemental file.

## Acknowledgements

We thank Dr. Ananya Chatterjee for the construction of multiple cloning site (MCS) containing pTurboGFP-B construct.

