## Supplementary for "An unusually long 5’UTR of ATP dependent cold shock DEAD-box RNA helicase gene *csdA* negatively regulates its own expression in *Escherichia coli*"

Supplementary figure 1:

A.

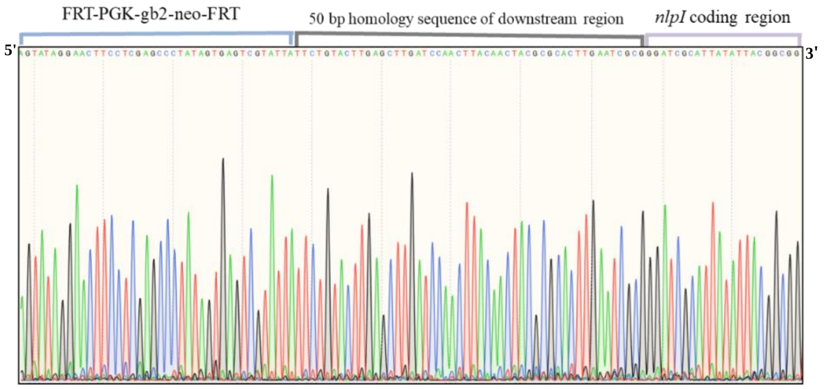


B.

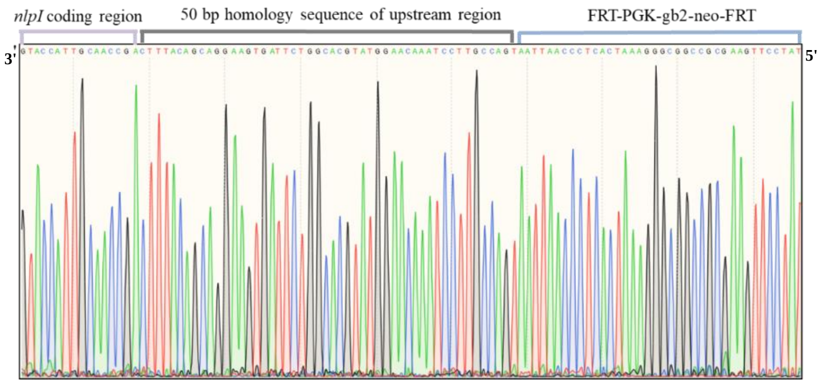


Fig. 1. (A) Sequence confirmation of the correct insertion inside the *E. coli* chromosome by using a forward primer (5' TATCAGGACATAGCGTTGGCTACC 3') which was complementary to a region located on the FRT-PGK-gb2-neo-FRT cassette, upstream of the 50 bases homology region at the 3' end of the 200 bases *csdA* promoter fragment. (B) A reverse primer was used (5' CGAGACTAGTGAGACGTGCTAC 3') which was also complementary to a region located on the FRT-PGK-gb2-neo-FRT cassette but downstream of the 50 bases homology region at the 5' end of the 200 bases *csdA* promoter fragment.

Supplementary figure 2:

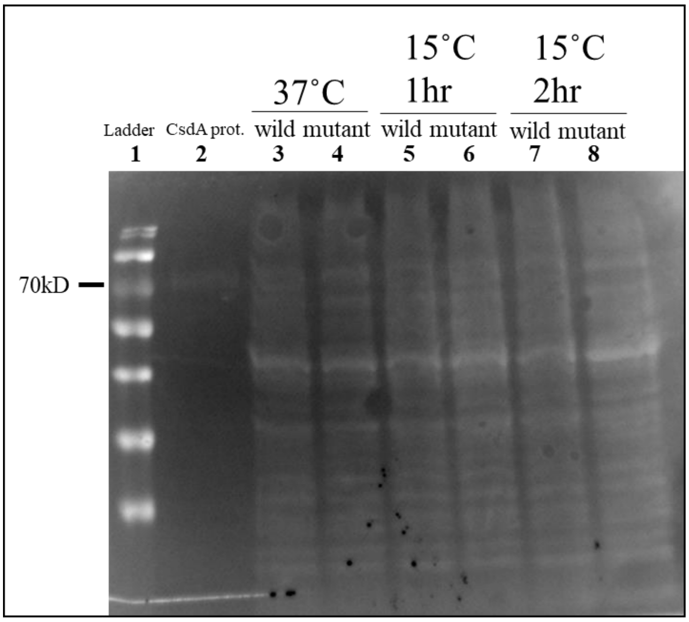


Fig. 2. Loading control: Equal amount of the total cellular proteins loaded which are stained with Ponceau S stain.
